# Differential expression of starch and sucrose metabolic genes linked to varying biomass yield in *Miscanthus* hybrids

**DOI:** 10.1101/2020.08.04.236885

**Authors:** Jose J De Vega, Ned Peel, Sarah J Purdy, Sarah Hawkins, Iain Donnison, Sarah Dyer, Kerrie Farrar

## Abstract

*Miscanthus* is a commercial lignocellulosic biomass crop owing to its high biomass productivity and low chemical input requirements. Interspecific *Miscanthus* hybrids with high biomass yield were shown to have low concentrations of starch and sucrose but high concentrations of fructose. We performed a transcriptional RNA-seq analysis between selected *Miscanthus* hybrids with contrasting values for these phenotypes to clarify how these phenotypes are genetically controlled. We observed that genes directly involved in the synthesis and degradation of starch and sucrose were down-regulated in high yielding *Miscanthus* hybrids. At the same time, glycolysis and export of triose phosphates were up-regulated in high yielding *Miscanthus* hybrids. Our results evidence a direct relationship between high expression of essential enzymatic genes in the starch and sucrose pathways, high starch concentrations, and lower biomass production. The strong interconnectivity between genotype, chemotype and agronomic traits opens the door to use the expression of well-characterised genes in the starch and sucrose pathway for the early selection of high biomass yielding genotypes from large *Miscanthus* populations.

## INTRODUCTION

*Miscanthus* is a candidate biofuel crop owing to its high biomass yield and low input requirements [1, 2]. It is also naturally adapted to a wide range of climate zones and land types [3, 4]. Currently, *Miscanthus* is mainly used for combustion, but there is keen interest in its development as a sustainable substrate for bioethanol or biomethane production [5, 6].

*Miscanthus* is a C4 perennial rhizomatous grass crop closely related to sugarcane (*Saccharum* spp.), sorghum (*S. bicolor*) and maize (*Zea mays*). However, unlike these species, *Miscanthus* is a non-food crop and can be grown on lower agricultural grade or marginal land so as not to compete with food production [7, 8].

Natural interspecific hybridisation events occur between several *Miscanthus* species with overlapping geographic distributions [9]. The main commercial *Miscanthus* genotype to date, *M. x giganteus*, is a sterile triploid wild hybrid resulting from the hybridisation between a diploid *M. sinensis* and a tetraploid *M. sacchariflorus. M. x giganteus* has desirable traits, including high yield and early establishment [10-12]. However, *M. x giganteus* must be clonally propagated, which doubles establishments costs compared to a seed-based option [13]. Therefore, several European breeding programmes are aiming to develop a seed-based crop through recreating the hybridisation event between *M. sinensis* and *M. sacchariflorus* to produce new hybrids that out-perform *M. x giganteus* [13, 14]. The hybrids produced at IBERS (Wales, UK) exhibited strong heterosis for several traits and have been characterised in previous publications [15-18].

A major hindrance to the improvement of perennial energy crops through breeding is the long duration for new crosses to reach the maturity stage when their traits can be assessed [18]. This creates a pressing need for the identification of molecular markers to predict yield before maturity is reached. Genetic markers were identified via association mapping for seven-teen traits in *Miscanthus* [19], and metabolic biomarkers successfully predicted the final yield eight months later [17]. The identification of transcriptional predictors in *Miscanthus* could provide a cost-effective tool to accelerate selection either using expression level as a marker or by identifying new target genes.

*Miscanthus* is harvested for the structural cell-wall polysaccharides, and as a result, multiple studies have focused on its structural carbohydrates [20, 21]. However, it is the processing and storage of non-structural carbohydrates (NSC), such as sucrose and starch, that underpin biomass traits [17].

We have previously shown that high yielding *Miscanthus* genotypes from an interspecific hybrid mapping family had low starch concentrations in the stem and a low ratio of starch to fructose [17]. These distinctive carbohydrate profiles were consistent across years and growing environments; thus, the phenotype is likely to be genetically controlled [17, 22]. Unlike many C3 temperate grasses, C4 species such as *Miscanthus* or maize do not accumulate fructans but instead accumulate starch as a transient form of storage carbohydrate [23, 24]. The concentration of starch in the mapping family was up to 15 % of the dry weight (DW) on average. However, higher values were observed in the lowest yielding lines, raising the possibility to bred “starch-cane” *Miscanthus* for liquid biofuel or biogas generation [25]. Identifying differentially expressed genes (DEGs) that relate to the carbohydrate profile could further facilitate breeding for such traits.

In this study, we analysed root, stem and leaf RNA-seq data from the hybrid progeny from a cross of a diploid *M. sacchariflorus* genotype and a diploid *M. sinensis* genotype, which had contrasting carbohydrate profiles and yield measurements. We identified differentially expressed genes associated with the observed metabolic profiles, using the recently completed *M. sinensis* reference genome (*Miscanthus sinensis* v7.1 DOE-JGI). Integrating expression and metabolic data is a logical strategy given the strong interconnectivity between genotype, chemotype and phenotype, and the lower genetic complexity of intermediate phenotypes, such as metabolites and yield subcomponents [26, 27].

## METHODS

### Mapping population establishment and phenotyping

A total of 102 genotypes from a paired cross between diploid *M. sinensis* genotype “*M. sinen* 102” and a diploid *M. sacchariflorus* genotype “*M. sacch* 297” were sown from seed in trays in a glasshouse in 2009. In 2010, individual plants were split to form three replicates of each genotype and then planted out into the field in a spaced-plant randomised block design comprising three replicate blocks at IBERS, Aberystwyth, UK. Details of the phenotyping were previously described [17]. Briefly, the family was harvested in February 2015 following the 2014 growing season. Biomass was dried to a constant weight, and the average DW weight per plant (kg) was calculated. Soluble sugars were extracted and quantified enzymatically and photometrically from known standard curves on the same plate, as previously detailed [18]. Starch was extracted using a modified *Megazyme* commercial assay procedure and quantified photometrically from known standard curves on the same plate, as previously described [18]. Four hybrid genotypes were selected based on a low or high number of tillers (transect count of tillers). Correlation between concentrations, plant height and biomass phenotypes for the whole mapping population was previously quantified [17]. Pearson’s correlation values between the number of tillers and the other phenotypes were determined for the whole family. Differences between the four selected hybrids for all phenotypes were determined with Student’s two-tailed t-tests.

### RNA sequencing and pre-processing

RNA was extracted from the four selected hybrids, as well as from the two parents of the family. Extraction was performed using RNeasy Plant Mini kit (Qiagen, CA, USA) according to the manufacturer’s instructions. Total RNA samples were sent to the sequencing service at the Earlham Institute (Norwich, UK) where standard Illumina RNA-seq libraries were prepared and sequenced using the HiSeq 2000 platform. The raw reads were filtered with Trim Galore [28] using the default options for paired-end reads to remove Illumina adaptor sequences and reads with quality scores below 20. Cleaned reads were aligned to the *M. sinensis* reference genome (*Miscanthus sinensis* v7.1 DOE-JGI, http://phytozome.jgi.doe.gov) downloaded from Phytozome with STAR using the “2-pass” mode [29] and Kallisto using the “quant” mode with default options [30]. In both cases, the reference was indexed using the *M. sinensis* gene annotation (*Miscanthus sinensis* v7.1 DOE-JGI, http://phytozome.jgi.doe.gov) downloaded from Phytozome in GFF3 format. This same gene annotation was functionally annotated with GO terms and enzyme codes with the command-line version of Blast2GO [31] using BLASTX with an E-value of 1e-10 and the NCBI non-redundant (nr) and EBI InterPro databases.

### Differential expression and enrichment analysis

The differential expression and enrichment analysis are fully available in an R notebook (See Data availability), which is also available via Github. Briefly, Kallisto count files, one from each of the 23 libraries, were imported in R using TXimport [32]. Differential analysis was performed using DESeq2 [33] for each tissue (root, stem, leaf) independently. Raw gene counts were obtained from Kallisto alignments and normalised using DESeq2 for the top 1,000 most variable genes to cluster the samples. Genes with a False Discovery Rate (FDR) < 0.05 were considered differentially expressed (DE). We compared two groups of hybrids; each hybrid group was composed of two genotypes (genotypes 112 and 90 against genotypes 18 and 120). We also compared the hybrids against the *M. sacchariflorus* and *M. sinensis* parent, one at the time. A gene only was considered DE between hybrids and parents when it was DE against both parents. The enrichment analysis was based on an F-fisher test (FDR▫< ▫0.05) using the library topGO [34] with the “weight01” algorithm. Using the lists of DE genes and functional annotation as inputs, topGO compared the number of DEGs in each category with the expected number of genes for the whole transcriptome. The “weight01” algorithm resolves the relations between related GO ontology terms at different levels. Enriched categories were plotted using ggplot2 [35]. Genes in enriched GO terms were further analysed using the online Phytomine [36] and Thalemine [37] databases. Genes annotated with enzyme codes were plotted using the online KEGG mapper [38].

## RESULTS

### Contrasting carbohydrate metabolism in sequenced genotypes from a *Miscanthus* mapping family

A total of 102 genotypes from a paired cross between diploid *M. sinensis* (“*M. sinen* 102”) and a diploid *M. sacchariflorus* (“*M. sacch* 297”) were established in field conditions and phenotyped. Non-structural carbohydrates were sampled in July 2014, during the summer growing season, and annual yield was obtained at harvest after the following winter. The distribution of carbohydrate concentrations and biomass yield for 98 hybrids were previously reported [17]. After including additional information about number of tillers for the population (Fig. 1 and Suppl. Table S1), we observed significant correlations between number of tillers and starch (r = -0.45, p < 0.001), fructose (r = 0.31, p < 0.005), and total NSC (r = -0.40, p < 0.0001) for the whole family (Suppl. Table S1). We also observed significant correlations between number of tillers and the ratio of sucrose/starch (*r* = 0.37, p < 0.001), fructose/starch (r = -0.45, p < 0.001), glucose/starch (r = -0.38, p < 0.001) and sucrose/fructose (r = -0.32, p < 0.01). We observed a significant positive correlation between biomass yield and number of tillers (r = 0.62 ± 0.03 for three seasons, p < 0.001).

**Figure 1:**
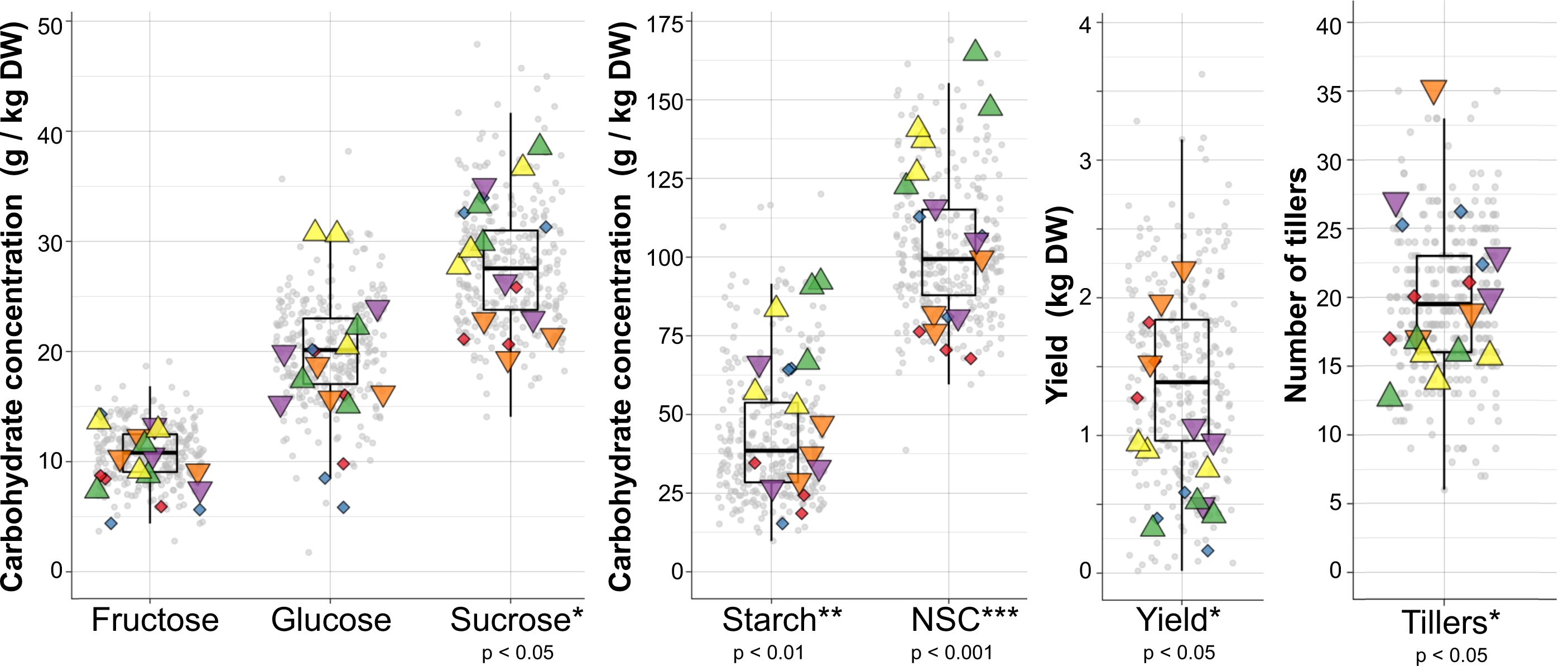
Concentrations of non-structural carbohydrates, number of tillers, and biomass yield in a mapping population comprised of 102 *M. sinensis* X *M. sacchariflorus* hybrids. Values for four hybrids with contrasting phenotypes (“high” and “low”), which were selected for RNA sequencing, are highlighted (Triangles). Significant differences (T-test) between the hybrids are annotated under each phenotype. The two parents of the family were also sequenced and phenotyped (diamonds). Boxplots summarise the distribution of values for the whole family for each phenotype.

Four *M. sinensis* X *M. sacchariflorus* hybrids from this family (Triangles in Figure 1) were selected for RNA sequencing in their fourth growing season (2013), based on a higher or lower than the average number of tillers. The two parents of the family were also sequenced (Diamonds in Figure 1). When the four sequenced hybrids were divided into two groups (genotypes 112 and 90 against genotypes 18 and 120), we observed significant differences between these groups in the number of tillers (p < 0.05), biomass yield quantified as dry weight per plant (p < 0.05), and the final canopy heights (p < 0.05). We also observed a significant difference between these two groups in the concentrations of starch (p < 0.005) and sucrose (p < 0.05), but we did not observe significant differences between groups in the concentrations of fructose or glucose. The most significant difference (p < 0.001) was observed in the total concentration of non-structural carbohydrates (NSC), which was calculated as the sum of the glucose, fructose, sucrose and starch concentrations. We observed significant differences also in the fructose/starch (p < 0.05) and glucose/fructose ratios (p < 0.01). However, any other ratio between concentrations was not significantly different between the groups (Suppl. Table S2).

There was a significant difference between the *M. sacchariflorus* and *M. sinensis* parents in NSC (p < 0.05) and sucrose concentrations (p < 0.01). However, there was no significant difference between the parents in the starch, fructose or glucose concentrations (Suppl. Table S2). It is likely an example of heterosis (transgressive segregation) that significant differences in starch, fructose or glucose concentrations were observed in the hybrid progeny but not the parents.

### Differential expression (DE) analysis between hybrids and species

We performed RNA-seq from the leaf, stem and root tissue samples extracted from four *M. sacchariflorus* X *M. sinensis* interspecific hybrids, and their two parents (Table 1). When the normalised counts obtained from DESeq2 [33] were used to cluster the samples (Figure 2), the samples firstly grouped by tissue (PC1) and secondly by species (PC2). As a result, the downstream analysis was performed for each tissue separately. Stem and root samples clustered together, and the clustering of these separately from the leaf tissue explained 64% of the variation. Species explains 17 % of the variation, with the hybrids falling between the two parents, which were furthest apart from each other.

**Table 1:** RNA-seq libraries used in this study.

| Genotype | tissue | Library | Group |
| --- | --- | --- | --- |
| 112 (hybrid) | root | LIB2338 | High NSC / Low<br>yield / Fewer<br>tillers |
|  | stem | LIB2339 |  |
|  | leaf | LIB2340 |  |
| 90 (hybrid) | root | LIB2341 |  |
|  | stem | LIB2342 |  |
|  | leaf | LIB2343 |  |
| 120 (hybrid) | root | LIB2344 | Low NSC / High<br>yield / Many<br>tillers |
|  | stem | LIB2345 |  |
|  | leaf | LIB2346 |  |
| 18 (hybrid) | root | LIB2347 |  |
|  | stem | LIB2348 |  |
|  | leaf | LIB2349 |  |
| *M. sacch* 297 | stem | LIB2352 | Progenitors |
|  | leaf | LIB2353 |  |
|  | stem | SAM1158 |  |
|  | root | SAM1159 |  |
|  | leaf | SAM1160 |  |
|  | root* | SAM1161 |  |
| *M. sinensis* 102 | stem | LIB2350 |  |
|  | leaf | LIB2351 |  |
|  | stem | SAM1162 |  |
|  | root | SAM1163 |  |
|  | leaf | SAM1164 |  |

*M. sacch* = *M. sacchariflorus*; \*root tip

**Figure 2:**
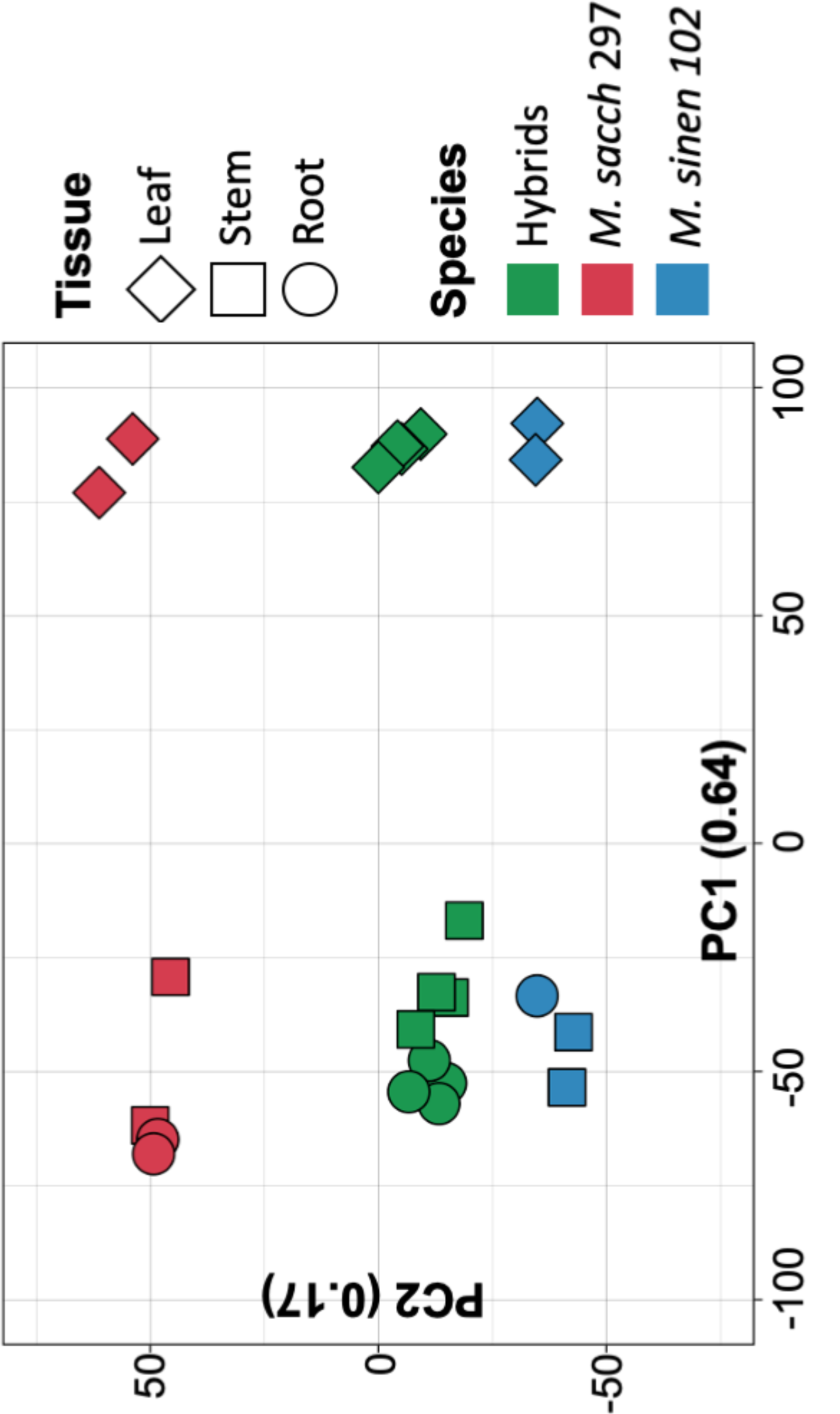
Principal component analysis of the normalised gene counts from 23 RNA-seq libraries generated from leaves (diamonds), stems (squares) and roots (circles) obtained from four *M. sinensis* X *M. sacchariflorus* hybrids (green shapes) with contrasting phenotypes and their parents (red and blue shapes). Gene counts were obtained from Kallisto alignments and normalised using DESeq2 for the top 1,000 most variable genes.

We obtained 1,386 differentially expressed genes (DEG; Suppl. Table S3) in total between the hybrids identified as “High NSC” and “Low NSC” (Figure 1) at FDR < 0.05 (Figure 3A). There were 892 DEGs in stems (598 up-regulated and 294 down-regulated), 741 DEGs in leaves (410 up-regulated and 331 down-regulated), and only 253 DEGs in roots (116 up-regulated and 137 down-regulated). 64 % of the DEGs in roots were DE in both of the other tissues too, but most DEGs in stem or leaves were exclusively DE in either stem or leaves.

**Figure 3:**
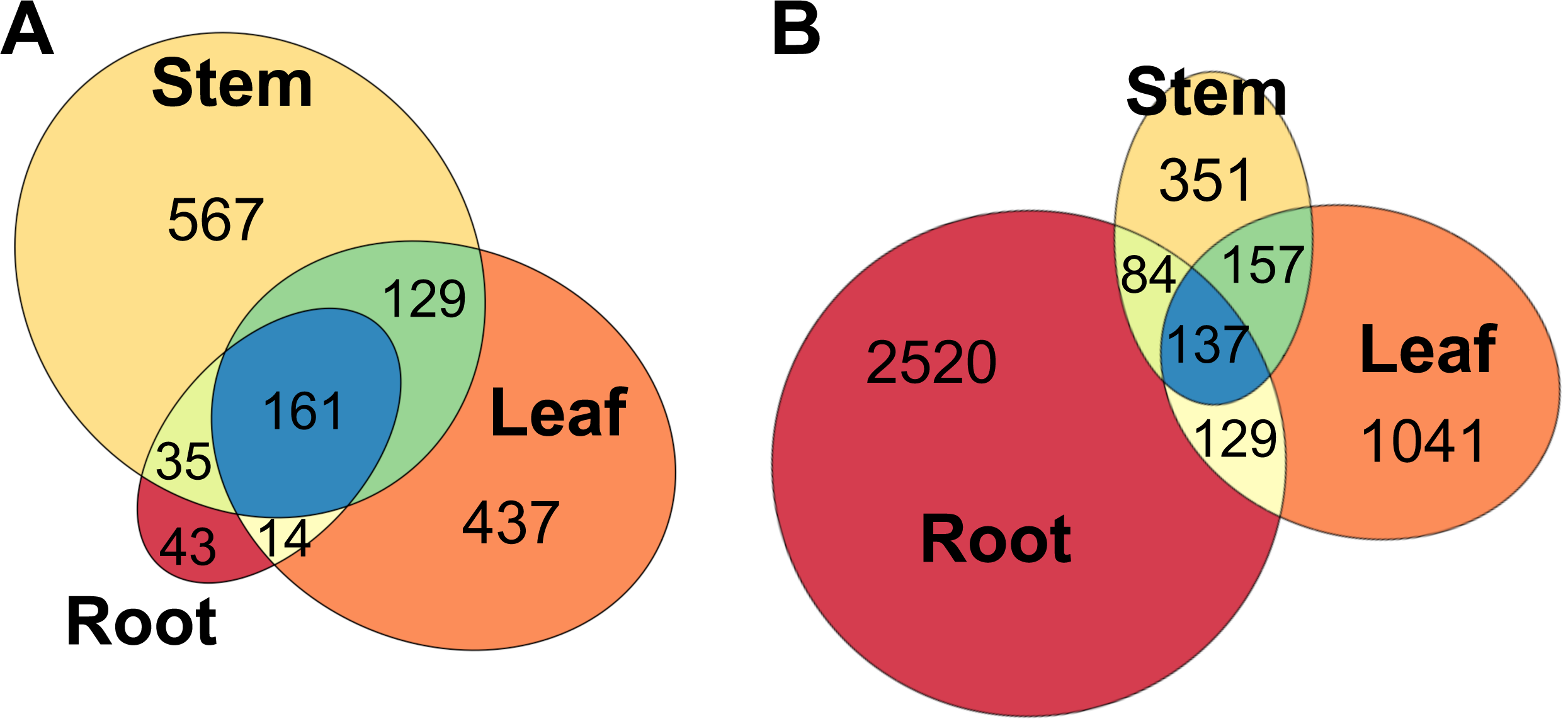
Number of differentially expressed genes shared between root, leaf and stem tissues among the “High NCS” and “Low NCS” *Miscanthus* hybrids at FDR < 0.05 (A), and between the hybrids and their progenitors (B). A gene only was considered DE between hybrids and parents when it was DE against both parents.

We also compared the expression between the hybrids against each parent and considered a gene as DE if it was DE in both comparisons at FDR < 0.05 (Suppl. Table S4). Under these criteria, there were 2,870 DEGs in roots, 1,464 DEGs in leaves, and 729 DEGs in stems (Figure 3B). Only 64 among these DEG were also DE between “High NSC” and “Low NSC” hybrids. There were 16,311 DEGs between the hybrids and *M. sinensis* alone (Suppl. figure S1), and 15,616 DEGs between the hybrids and *M. sacchariflorus* alone (Suppl. figure S2), this is over a third of the total transcriptome.

### Enriched Gene Ontology (GO) terms in DEGs

Enrichment analysis of GO terms over-represented among DE genes allowed us to identify the biological processes (BP) and molecular functions (MF) that are differentially regulated in each tissue. After annotating the reference transcriptome with the homologous proteins and full set of GO terms and (Suppl. Table S5), we simplified the results to the more general “GO slim” terms.

All the significant enrichment “GO slim” terms among DEGs between the “High NSC” and “Low NSC” hybrids were associated with metabolic processes, with the single exception of “response to stress” in stems (Figure 4; Suppl. Table S6). Among the GO terms in the “biological process” category, the most significantly enriched ones (p < 0.001) were “Carbohydrate metabolism” and “Secondary metabolism” in stem and leaves, and “Generation of precursor metabolites and energy” and “response to stress” in stems. Among the “molecular process” category, “hydrolysis on glycosyl bonds” and “redox activities” were the most significantly enriched (p < 0.0001) in both stems and leaves (Suppl. Table S6).

**Figure 4:**
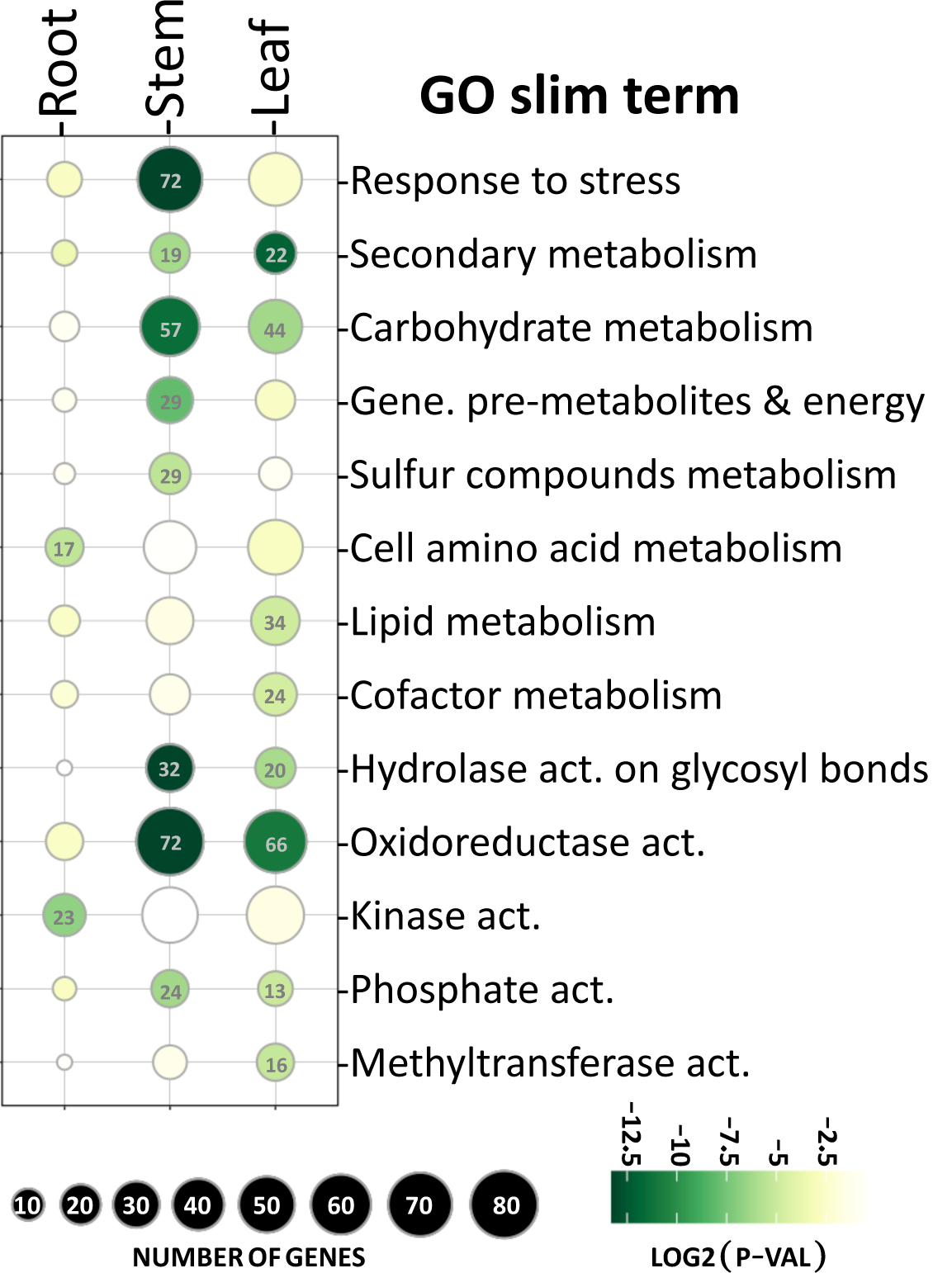
GO SLIM terms (rows) that were significantly enriched (p < 0.05) in each tissue (columns) among differentially expressed genes (DEG) from the “High NCS” and “Low NCS” *Miscanthus* hybrids DE analysis. The size of a bubble is proportional to the number of DEG annotated with that GO term. Rows are sorted by descending p-value (F-fisher test) and the bubble colour is representative to the obtained p-value, from lower (dark green) to higher (light green). Yellow (p > 0.05) and white (p > 0.1) bubbles were not enriched. All the enriched GO SLIM terms for the “biological process” (top 8 rows) and “molecular function” (bottom 5 rows) GO categories were included.

Thirty-six enzymatic reactions were annotated among DEG in the stem (Table 2). Only six were down-regulated in “High NSC”; four involved in the generation of precursor metabolites and energy, namely 6-phosphofructokinase (EC 2.7.1.11) and Triose-phosphate isomerase (EC 5.3.1.1) in the glycolysis pathway; Malate dehydrogenase NADP(+) (EC 1.1.1.82) in the pyruvate metabolism; and 2-carboxy-D-arabinitol-1-phosphatase (EC 3.1.3.63); and one each in the other GO categories, namely Beta-N-acetylhexosaminidase (EC 3.2.1.52) and carboxypeptidase (EC 3.4.16.-).

**Table 2:**
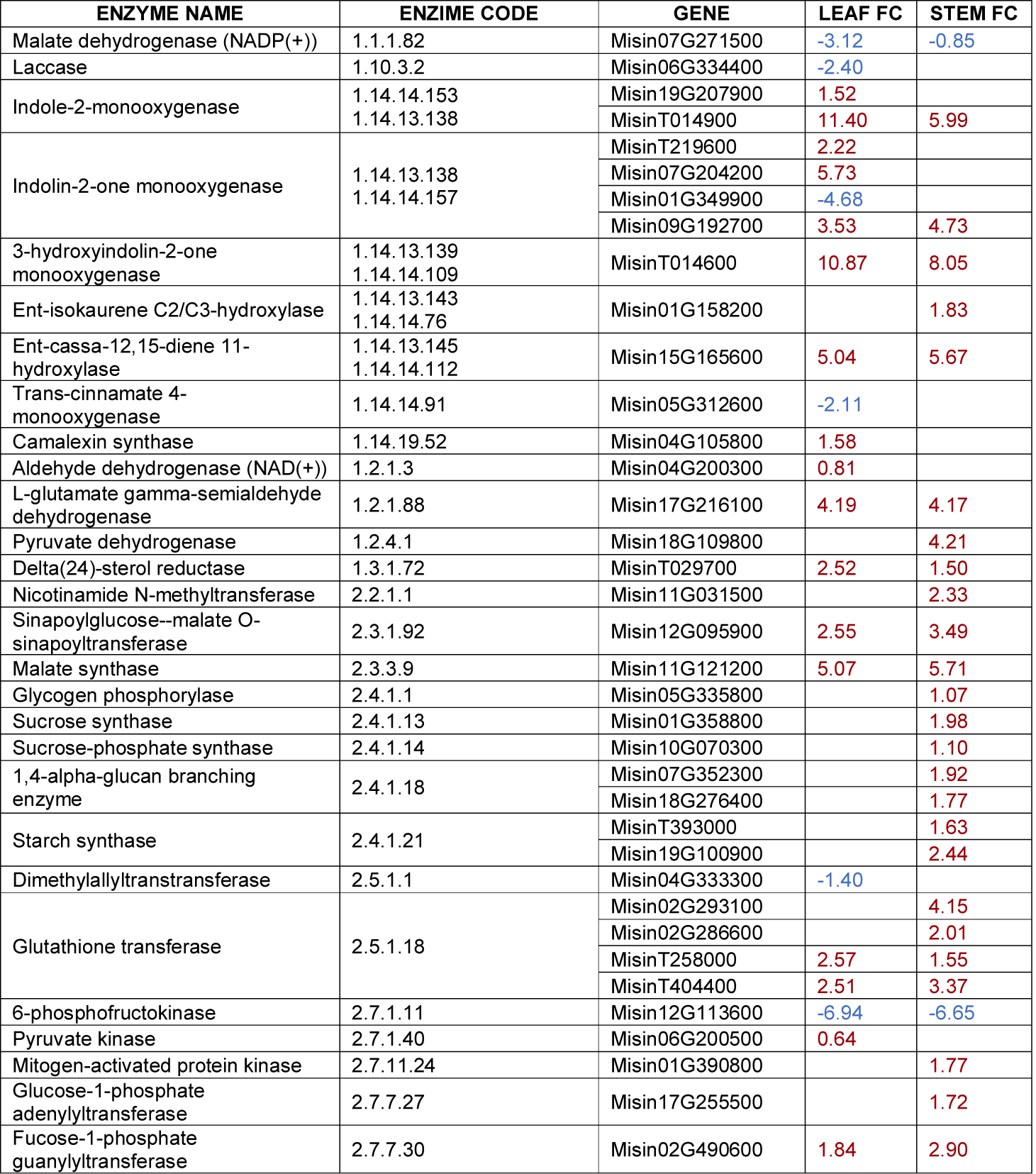
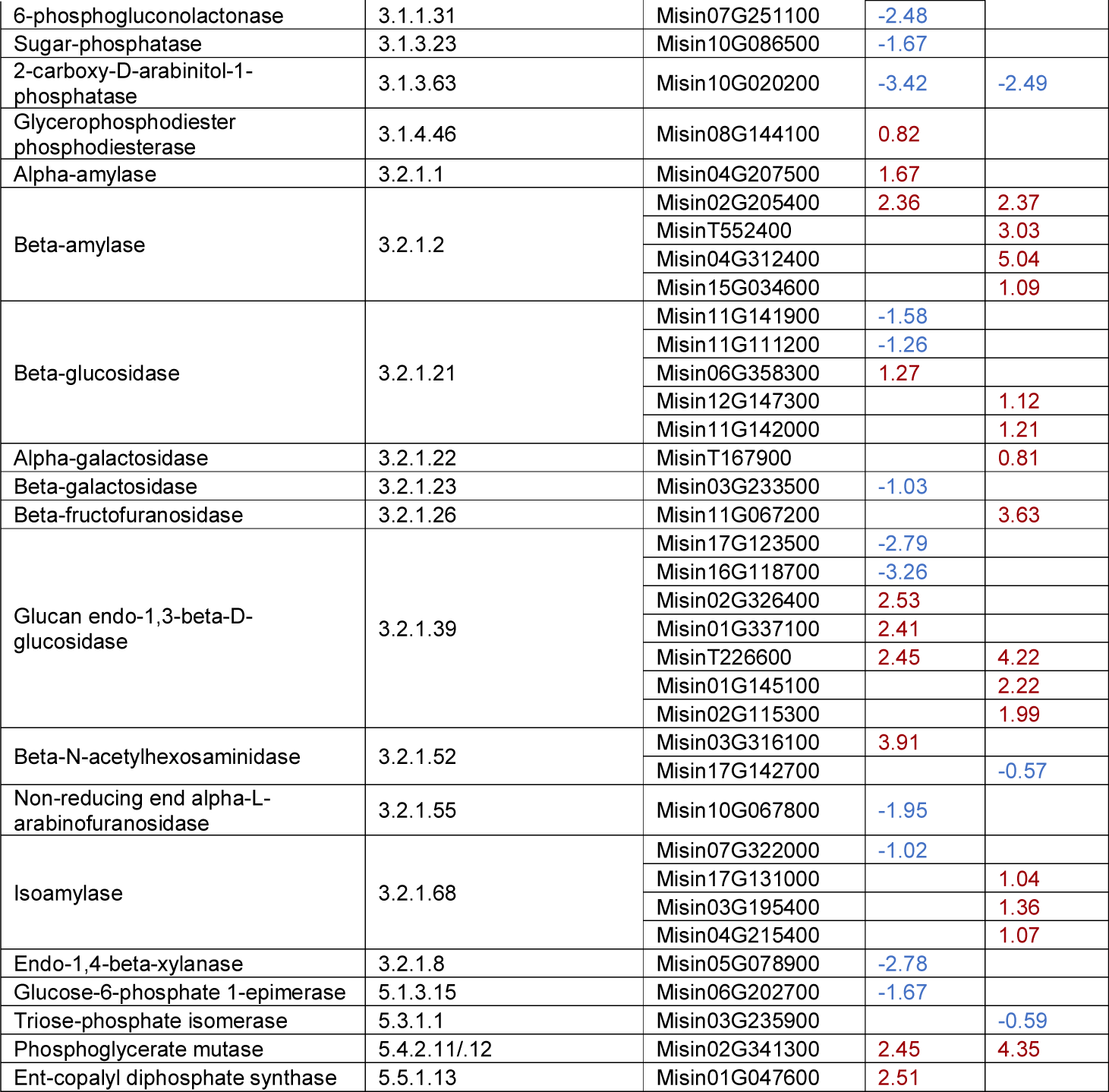
Carbohydrate and secondary metabolic enzymatic reactions differentially expressed between “high NSC” and “low NSC” *Miscanthus* hybrids. *Leaf FC or Stem FC = Log2 fold-change expression between “high NSC” / “Low NSC” hybrids in either leaf or stem tissues*.

A similar analysis on the enriched GO slim terms among DEGs between hybrids and parents (Suppl. Figure S3; Suppl. Table S7) revealed that the most significantly enriched GO terms (p < 0.01) were in the root and associated with RNA/DNA binding and translation (including ribosome biogenesis and equivalent terms), and several biosynthetic processes. Remarkably, there were no enriched GO terms in the stem between hybrids and parents.

### DEG associated with the starch and sucrose metabolism

There were 88 DEGs associated with the enriched “Carbohydrate metabolism” GO term (Suppl. Table S8), specifically 57 DEGs in stems (42 up-regulated and 15 down-regulated) and 44 DEGs in leaves (20 were up-regulated and 24 down-regulated). Thirteen DEGs were common to both tissues and showed close fold-change values in both tissues. All but two of these 88 DEGs could be functionally annotated, 52 and 56 of them had a homologous protein in *A. thaliana* or rice, respectively.

Twenty-nine DEGs were involved in enzymatic reactions that were part of the starch and sucrose metabolic pathways (KEGG pathway ath00500; Suppl. Figure S4). Among these, all 20 DEGs in stems were up-regulated in “High NSC”, but half of the DEGs in leaves (which were beta-glucosidases) were down-regulated in “High NSC”. Enzymatic proteins in the starch degradation pathway were DE in root and leaves (e.g. AMY3, ISA3, BAM1). At the same time, sucrose metabolism genes in the cytosol were only DE in stems (SUS3, SPS5). Similarly, reactions involving ADP-glucose were only DE in stems (e.g. AGP, SS2, SS3, SBE2).

Twenty-nine genes were annotated as involved in the “generation of precursor metabolites and energy” (Suppl. Table S8), 17 of which could be annotated with an enzymatic code (KEGG pathway ath00010; Suppl. Figure S5). Six genes were involved in starch metabolism (ISA3, DBE1, PFK2, SBE2, PHS2). The phosphofructokinase 2 (PFK2) is the only one clearly down-regulated in “High NSC”. Among the others, a malate synthase (MLS) and an aldehyde dehydrogenase 12A1 involved in siRNAs generation, and an Fts protease (FTSH6) in the chloroplast were all highly up-regulated (FC > 5) in “High NSC”. On the other hand, triosephosphate isomerase (TIM) was down-regulated in “High NSC”.

The relation between 32 DEGs involved in the twelve DE enzymatic reactions in starch and sucrose metabolism, plus three of the glycolysis reactions are summarised in Figure 5 and Table 3.

**Table 3:**
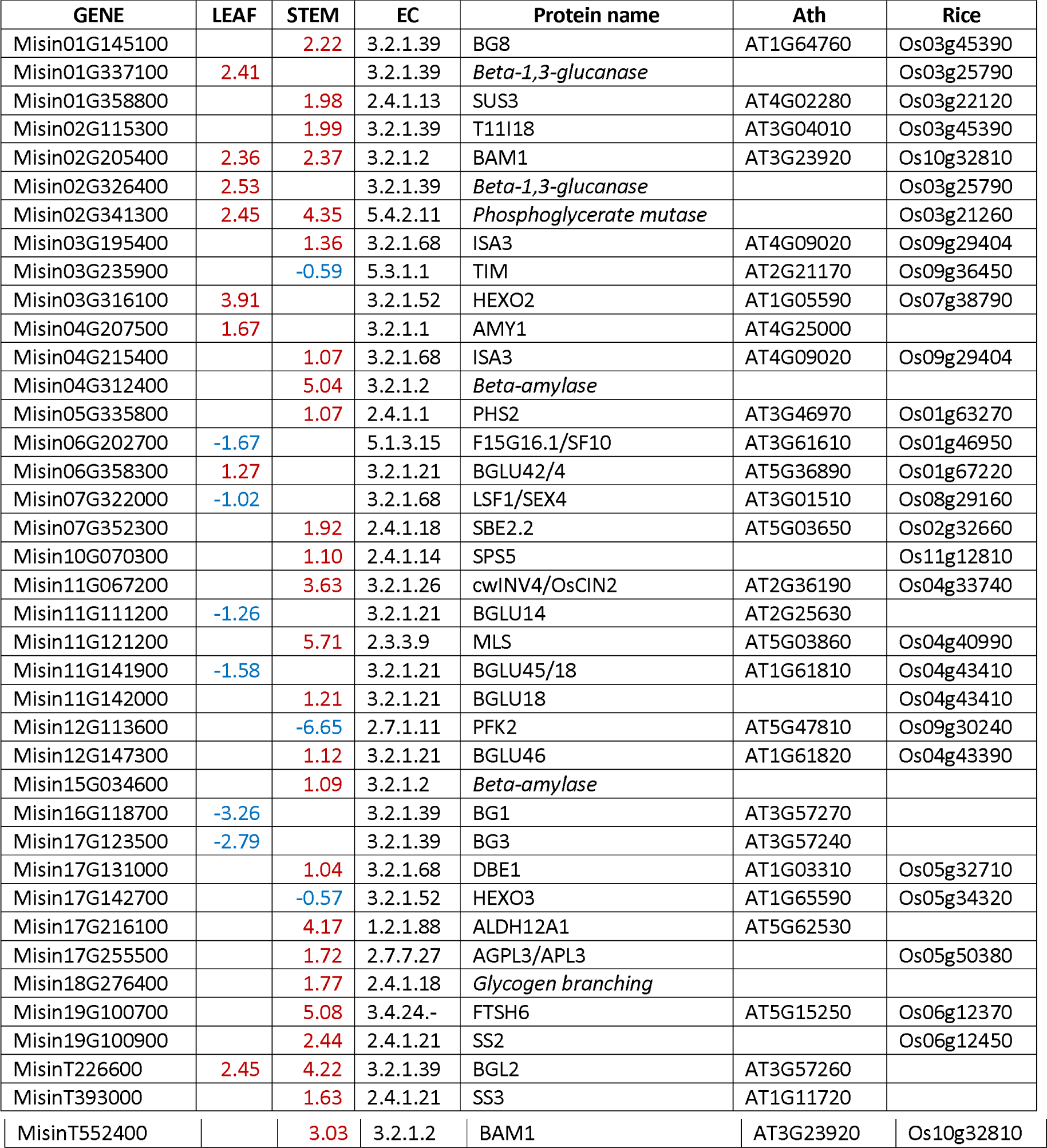
Thirty-nine differentially expressed genes were involved in twelve reactions in the starch and sucrose metabolism and three of the glycolysis reactions were highlighted in our analysis. *Leaf/stem = Log2 fold-change expression “high NSC” / “Low NSC” hybrids in either lead or stem tissues; Ath/Rice = Homologous protein in Arabidopsis thaliana and rice (The prefix “LOC_” is not included in the name)*.

**Figure 5:**
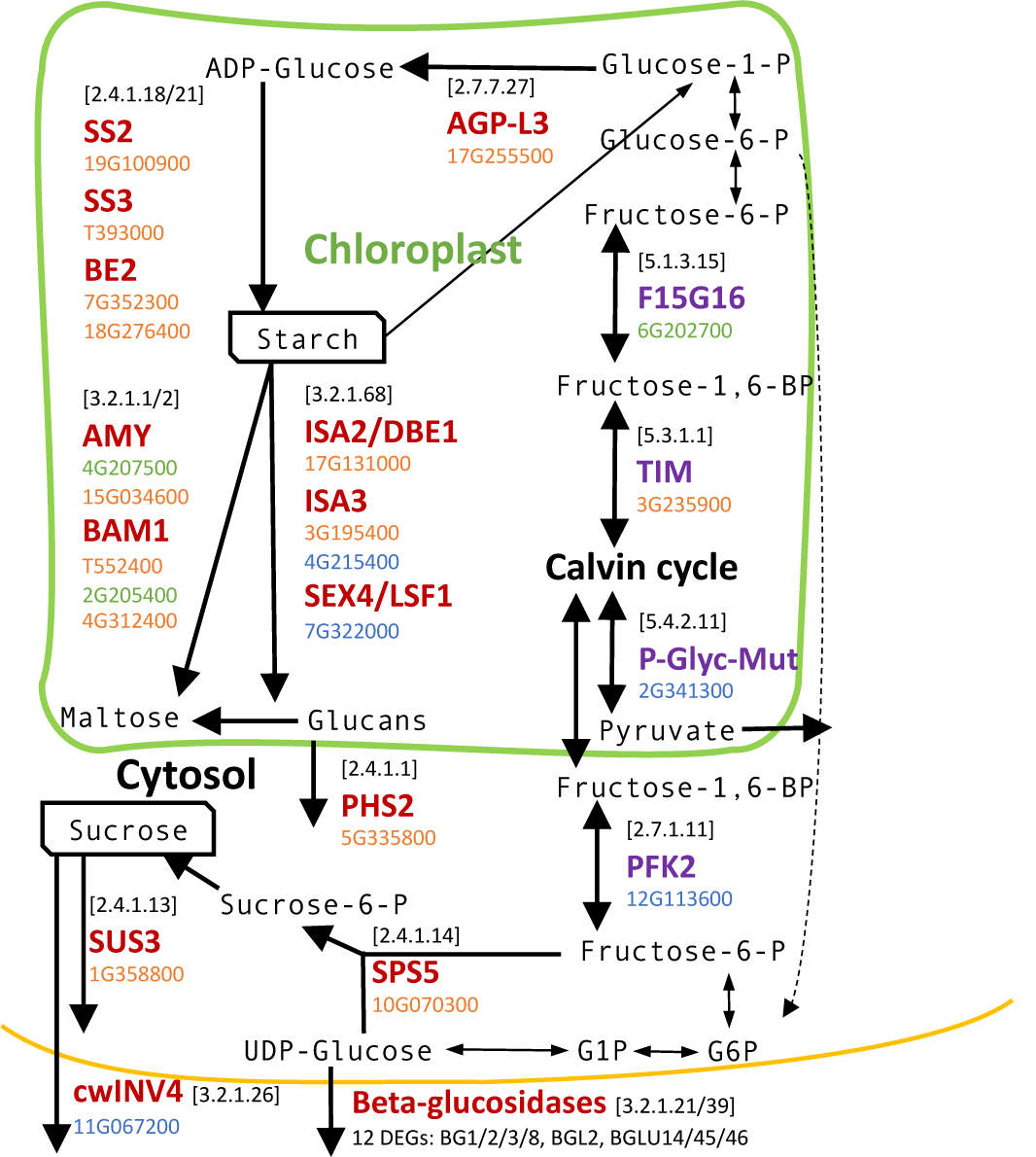
Schema of the starch and sucrose metabolism in plants, highlighting critical differentially expressed (DE) proteins between “High NSC” and “Low NSC” Enzymatic codes are shown between brackets. DE *Miscanthus* genes are included under their respective protein (The prefix “Misin_” is not included in the gene name). Genes were differentially expressed in leaves (coloured in green), stems (orange) or both tissues (blue).

### DEG associated with other enriched GO terms

The 72 genes annotated as “Response to stress” were involved in a broad range of responses (Suppl. Table S10). On the other hand, the most significantly enriched GO terms in the “Molecular functions” category were associated with metabolic-related enzymatic reactions, namely “oxidoreductase activities” and “hydrolase activities”. The former included 38 cytochrome P450 proteins.

“Secondary metabolism” was enriched in both stems and leaves. 17 of the 19 DEGs in stems were up-regulated, but half of the DEGs in leaves were down-regulated. 16 of the 31 genes involved in the “secondary metabolism” were cytochrome P450 proteins (Suppl. Table S11). Six were included in benzoxazinoids biosynthesis, which is associated with defence in grasses. Another six were involved in terpenoids and phenylpropanoid biosynthesis (KEGG ath00900 and ath00940).

Many of the identified DEG in enriched functions showed no homologies in model organisms and consequently remain uncharacterised. This is the case in 36 DE genes involved in the carbohydrate metabolism (over 88 total), whose function was evidenced by the presence of a protein domain, but with an unclear role. A similar case is noted in two genes involved in the “generation of precursor metabolites", twelve genes involved in the “secondary metabolism”, and 17 genes involved in “response to stress”.

## DISCUSSION

We performed a transcriptional RNA-seq analysis between selected *Miscanthus* hybrids with negative correlations between starch and sucrose concentrations and biomass yield.

Using a mapping family (n = 102) between a diploid *M. sinensis* and a diploid *M. sacchariflorus*, we previously demonstrated that high biomass yielding *Miscanthus* hybrids had low starch and high fructose concentrations in the stem, and a lower ratio of sucrose, glucose and starch to fructose under peak growing conditions [17]. Here, we selected four hybrids from this mapping family based on the number of tillers (transect count), which was shown to be an accurate predictive phenotype for biomass yield [15]. These four hybrids could be divided into two groups (Table 1), which showed significant differences in the concentrations of starch and sucrose, but not of hexose. The most significant differences were observed for total NSC because of the cumulative effect of the differences in starch and sucrose.

Approximately 10 % of the total genes were DE between these two groups of hybrids in stems and leaves, but not in roots. Among these DE genes, there was an enrichment of genes involved in carbohydrate and secondary metabolism in stem and leaves, and in the “generation of precursor metabolites and energy” in stem only. However, these GO terms for biological processes were not regulated similarly in both tissues. While the DEGs in the enriched categories were predominantly up-regulated in stems, they were evenly up-regulated and down-regulated in leaves. The DE of carbohydrate metabolising genes between the leaf, stem and root is to be expected as it has been previously reported that carbohydrates are differentially distributed between these tissues in *Miscanthus* in July, the same month our study was conducted [18, 22]. Specifically, the abundance of starch in stems was up to 6x more concentrated in the leaf than stem, the below-ground biomass preferentially accumulated starch, and soluble sugars tended to be in greater concentrations in the stems compared to leaves [22]. Our transcriptional observations therefore largely reflect the distribution of carbohydrates; starch metabolism transcripts were DE in the leaf where starch is the most abundant carbohydrate, and sucrose metabolising enzymes were DE in the predominantly sucrose accumulating stem [18]. Fewer DEGs were observed in roots. Seasonal carbohydrate profiling of rhizomes in four genotypes showed that the soluble sugar contents were similar between genotypes and across two sites located 340km apart [18].

We observed that multiple genes involved in the synthesis (AGP, SS2, SS3, BE2) and degradation of starch in the chloroplast (AMY3, ISA3, SEX4, BAM1) were down-regulated in high biomass yielding genotypes. We also observed down-regulation of genes involved in the synthesis (SPS5) and degradation (SUS) of sucrose in high biomass-yielding genotypes. Genes involved in the starch metabolic pathway are up-regulated by a high sugar status [39-41], as there was a negative relationship between yield and soluble sugar (i.e. high yielders had lower sugar), it is consistent that the expression of sugar stimulated genes would be lower in high yielding genotypes.

Contrary to this, we noticed the up-regulation with a high fold-change in high biomass yield genotypes of triosephosphate isomerase TIM/PDTPI, which encodes a plastidic triose phosphate isomerase [42], and Phosphofructokinase 2 (PFK2). PFK2 catalyses the penultimate step before usable energy is extracted from the phosphorylated products of photosynthesis. This enzyme is, therefore, a main control point of glycolysis. The observation that high biomass plants have low carbohydrates can seem counter-intuitive, but the rationale is highly logical; high biomass plants maximise growth at the expense of their carbon reserves [43], whereas slow-growing types accumulate their reserves. The upregulation of the PFK2 gene encoding a major glycolytic enzyme is suggestive of a more rapid metabolism of photosynthate to fuel growth in the high yielding types. In summary, starch and sucrose synthesis was down-regulated in high yielding *Miscanthus* hybrids, while glycolysis and export of triose phosphates was up-regulated in high yielding *Miscanthus* hybrids.

These results support that high yielding *Miscanthus* genotypes were more rapidly accumulating structural mass, likely cellulose via sucrose metabolism [44-46], at the expense of starch [17, 18, 47]. The latter is further supported by the significant differences in the fructose-to-starch (but not glucose-to-starch) ratio between high and low yielding hybrids [17], which was also observed between the sequenced hybrids. Fructose is an indicator of sucrose metabolism, because it is produced exclusively from the metabolism of sucrose by the action of sucrose synthases (SUS), while glucose is produced by the metabolism of both sucrose and starch [48, 49]. Furthermore, in a C13 labelling experiment, it was observed that a greater proportion of the labelled carbon was observed in the insoluble fraction (mainly comprising cellulose) of a rapidly growing *Miscanthus* genotype, whereas a greater proportion was partitioned into starch in a slower-growing type [17]. Our results, therefore, add to these previous observations with the addition of transcriptomic evidence of the relationship between carbon metabolism, partitioning and growth.

We observed a significant enrichment of “response to stress” genes in stems. However, a further analysis did not reveal more details, only 32 of the 72 DEG in this category had a homologous protein in *A. thaliana*, and 23 of these were annotated as “response to stimulus", i.e. several types of environmental stress. Changes in starch metabolism are linked to changes in source-sink carbon allocation for protection against environmental stresses [50], and may expose differences between groups with phenotypic differences. On the other hand, several of the genes involved in the starch and sucrose metabolism are confirmed redox-regulated enzymes (e.g. AGP, SS3, BE2, AMY3, ISA3, SEX4, and BAM1), which partially explains the enrichment of in the “oxidoreductase activity” (GO:55114). The homologous in *A. thaliana* of *Miscanthus* genes annotated in “oxidoreductase activity” were usually involved in metabolic processes (GO:44699).

Our results evidence a direct relationship between high expression of essential enzymatic genes in the starch and sucrose synthesis pathway, high starch concentrations, and lower biomass production. The strong interconnectivity between genotype, chemotype and agronomic traits opens the door to use the expression of well-characterised genes in the starch and sucrose pathway for the early selection of high biomass yielding genotypes from large *Miscanthus* populations.

## ACKNOWLEDGEMENTS

We would like to thank Dr Anne Maddison for her technical support in generating the carbohydrate data. This work was funded by the Biotechnology and Biological Sciences Council (BBSRC) in projects BBS/E/T/000PR9818 (JDV), BBS/E/W/10963A01A (KF) and BB/CSP1730/1 (KF). Sequencing was funded through the “Capacity and Capability Challenge” funding awarded to Earlham Institute’s *Genomic Pipelines* group by BBSRC.

## DATA AVAILABILITY

Raw reads are deposited in the Short Reads Archive (SRA) under Bioproject ID PRJNA639832. The R code used in the analysis is deposited in Zenodo (http://doi.org/10.5281/zenodo.3834007) and Github (https://github.com/joseja/miscanthus_starch_rnaseq).

## SUPPLEMENTARY MATERIALS

Suppl. Figure S1: Number of differentially expressed genes shared between root, leaf and stem tissues between the hybrids and the *M. sinensis* progenitor.

Suppl. Figure S2: Number of differentially expressed genes shared between root, leaf and stem tissues between the hybrids and the *M. sacchariflorus* progenitor.

Suppl. Figure S3: GO SLIM terms (rows) that were significantly enriched (p < 0.05) in each tissue (columns) among differentially expressed genes (DEG) from the expression analysis between the hybrids and both progenitors. The size of a bubble is proportional to the number of DEG annotated with that GO term. Rows are sorted by descending p-value (F-fisher test) and the bubble colour is representative to the obtained p-value, from lower (dark green) to higher (light green). Yellow (p > 0.05) and white (p > 0.1) bubbles were not enriched. All the enriched GO SLIM terms for the “biological process” (top 8 rows) and “molecular function” (bottom 5 rows) GO categories were included.

Suppl. Figure S4: Down-regulated enzymatic reactions in the “starch and sucrose metabolism” pathway from KEGG (KEGG pathway ath00500) that were down-regulated in “high NSC” hybrids, which had higher concentrations of starch and sucrose.

Suppl. Figure S5: Enzymatic reactions in the “glycolysis/gluconeogenesys” pathway from KEGG (KEGG pathway ath00010) that were down-regulated (red boxes) or up-regulated (green boxes) in “high NSC” hybrids, which had higher concentrations of starch and sucrose.

Suppl. Table S1: Individual trait scores and Person correlation between traits.

Suppl. Table S2: Traits significantly different (T-test) between the sequenced samples.

Suppl. Table S3: Normalised counts, expression fold-change and P-values for all the genes in roots, stem and leaf tissue between groups of hybrids.

Suppl. Table S4: Normalised counts, expression fold-change and P-values for all the genes in roots, stem and leaf tissue between hybrids and parents.

Suppl. Table S5: Functional annotation, GO and enzyme codes for all the genes in the reference genome.

Suppl. Table S6: Enriched GO terms among DEG between groups of hybrids.

Suppl. Table S7: Enriched GO terms among DEG between hybrids and parents.

Suppl. Table S8: Detailed functional annotation of 88 DEG within the enriched “carbohydrate metabolism” GO term.

Suppl. Table S9: Detailed functional annotation of 29 DEG within the enriched “generation of precursor metabolites and energy” GO term.

Suppl. Table S10: Detailed functional annotation of 72 DEG within the enriched “response to stress” GO term.

Suppl. Table S11: Detailed functional annotation of 31 DEG within the enriched “secondary metabolism” GO term.

## Notes

### Competing Interest Statement

The authors have declared no competing interest.

