## Supplementary figures and images for "Differential expression of starch and sucrose metabolic genes linked to varying biomass yield in *Miscanthus* hybrids"

### Supplementary Figure S1

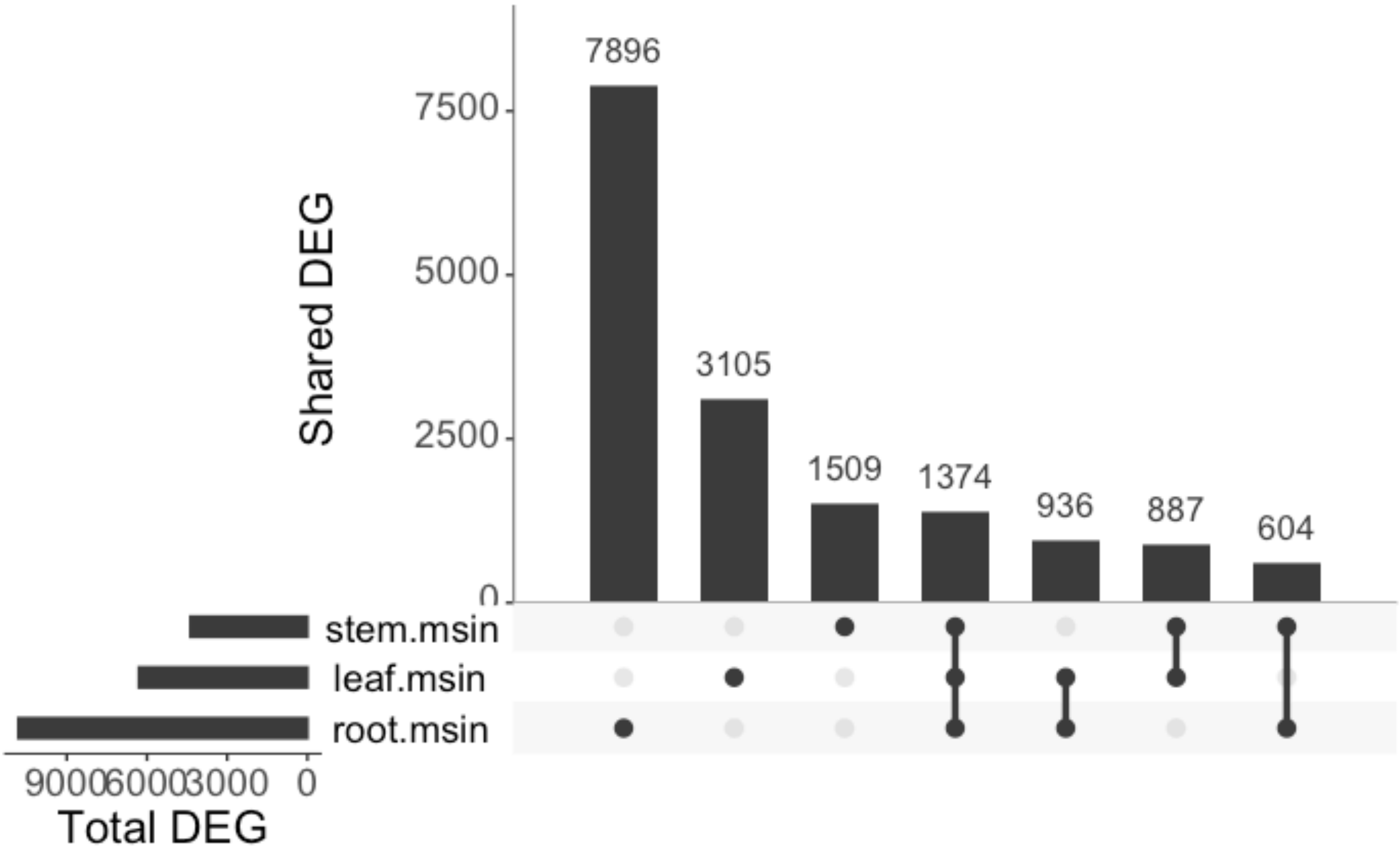

### Supplementary Figure S2

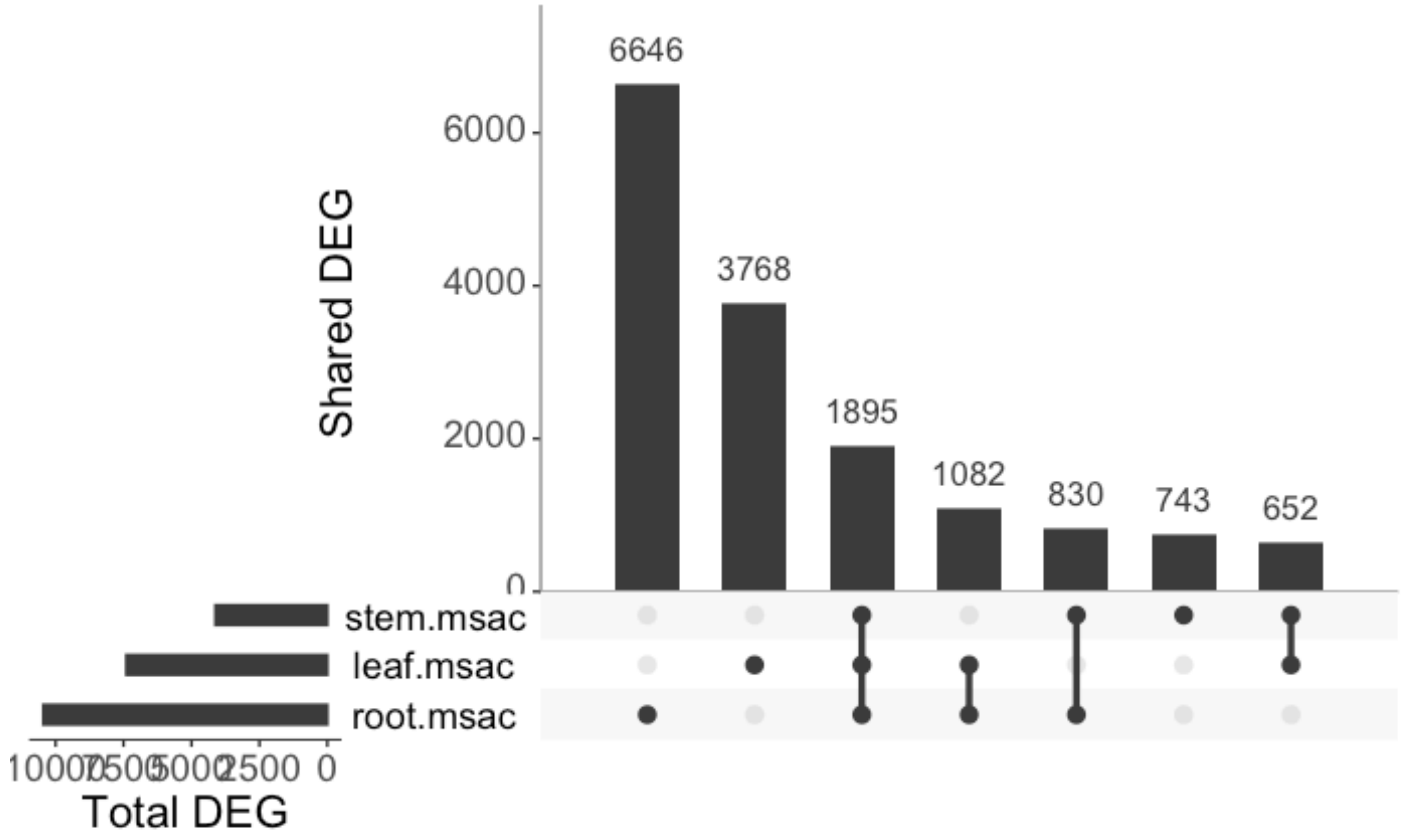

### Supplementary Figure S3

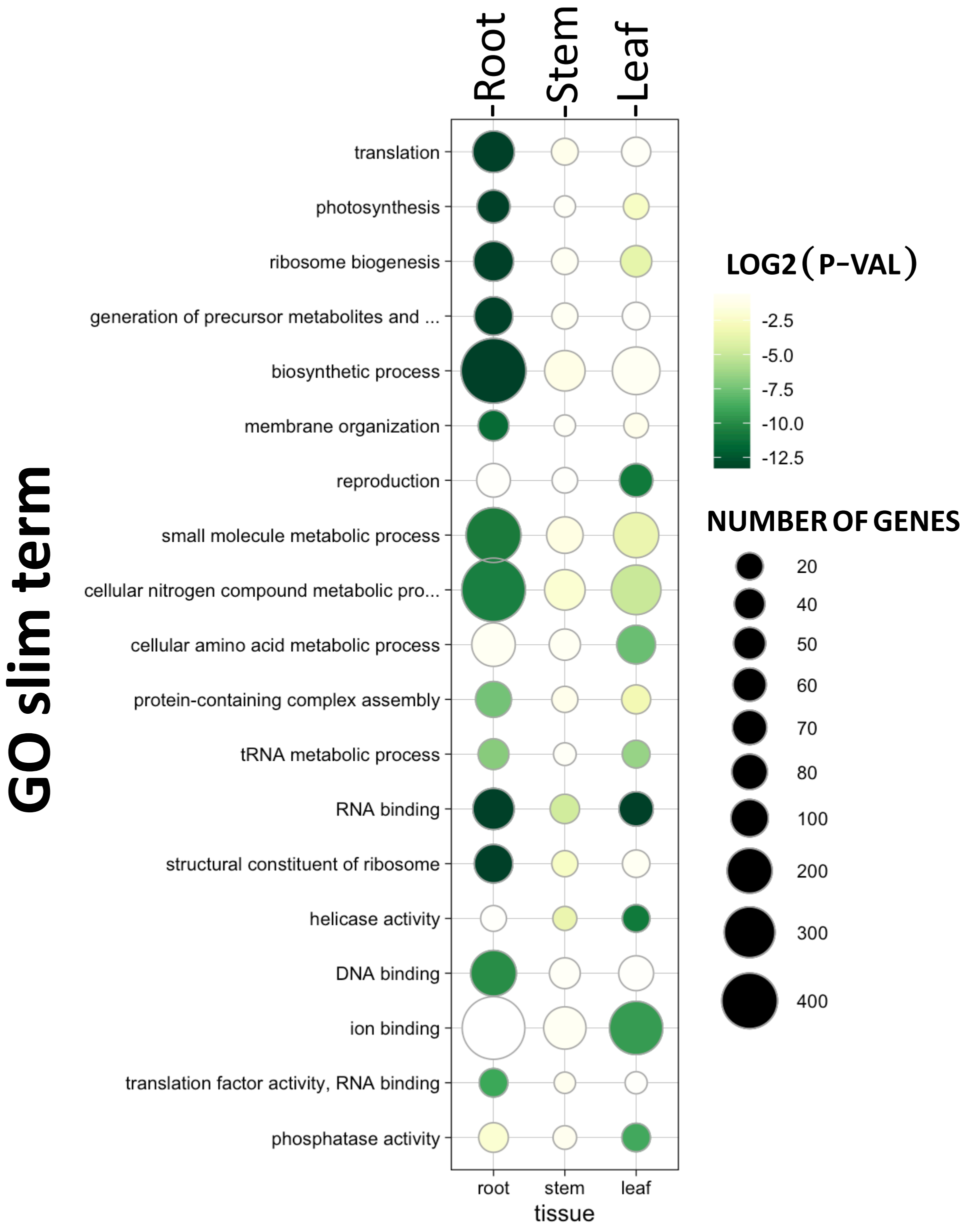

### Supplementary Figure S4

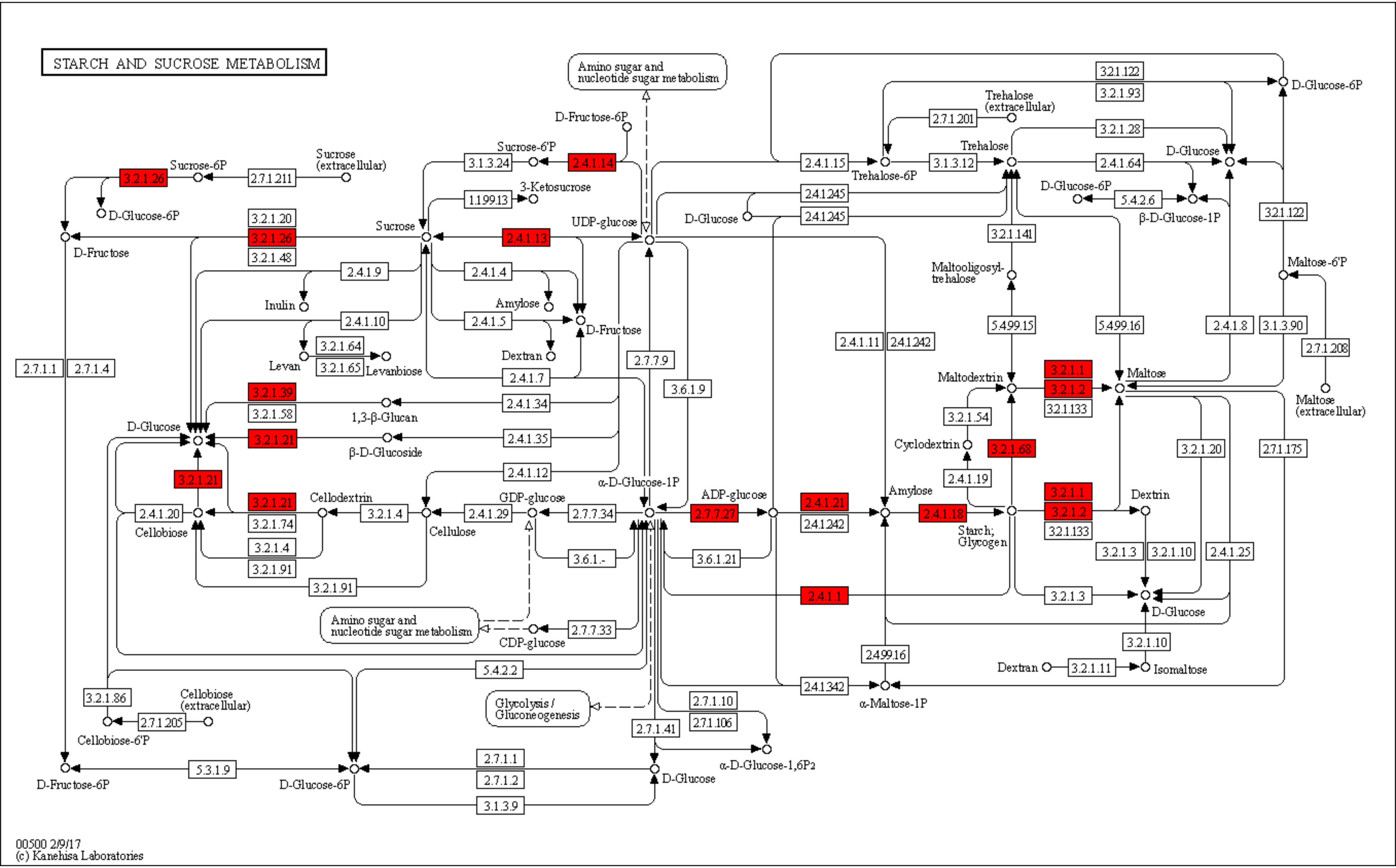

### Supplementary Figure S5

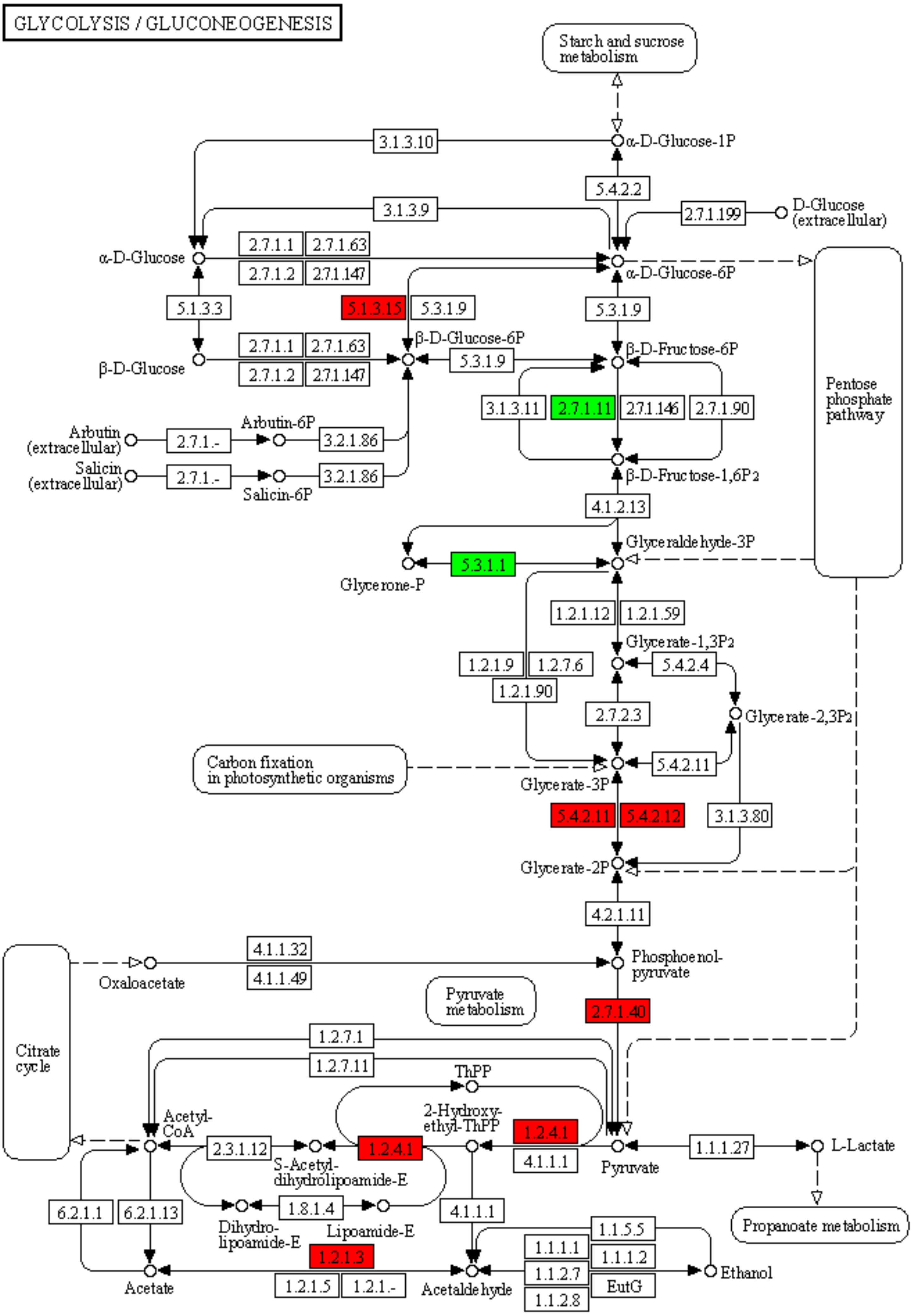


00010 4/12/18

(c) Kanehisa Laboratories
